# Purkinje Cell spike patterns do not correlate with nuclei cell spike patterns in mouse models for cerebellar disease

**DOI:** 10.1101/2025.05.19.654856

**Authors:** Alyssa M Lyon, Meike E van der Heijden

**Affiliations:** Graduate Program in Translation Biology Medicine and Health, Virginia Tech; Fralin Biomedical Research Institute, Virginia Tech Carilion; Center for Neurobiology Research, Virginia Tech Carilion; School of Neuroscience, Virginia Tech; Department of Pediatrics, Virginia Tech Carilion

## Abstract

Cerebellar dysfunction causes various movement disorders, including ataxia, dystonia, and tremor. Previous work demonstrated that spike patterns in cerebellar nuclei neurons were distinct between different movement disorder mouse models. However, often these changes arise from neural dysfunction in the cerebellar cortex, through misfiring, miswiring, or degenerating Purkinje cells. Even though Purkinje cells form the sole output from the cerebellar cortex, their information is relayed to other regions of the motor network via cerebellar nuclei cells. Purkinje cells make GABAergic synapses onto cerebellar nuclei cells, and it is often assumed that changes in Purkinje cell spike patterns result in inverse changes in nuclei cell spike patterns. Here, we test this hypothesis by answering the question of whether a reliable relationship between Purkinje cell and nuclei cell spike patterns exists. Single-cell, *in vivo* electrophysiology recordings of both cell types from six mouse models for cerebellar movement disorders were analyzed according to parameters relating to spike rate and irregularity. We investigated whether Purkinje cell spike patterns correlated with nuclei cell spike patterns. We found that some parameters for firing irregularity were positively correlated between Purkinje and nuclei cells but no – and particularly no inverse – relationship was observed between Purkinje and nuclei cell spike rate. Overall, this study begins to illuminate that the relationship between Purkinje cells and nuclei cell spike activity in a disease state is more complex and unpredictable. The data suggest Purkinje cell spike activity changes cannot accurately predict nuclei cell changes, which ultimately drive cerebellar disease states. Our findings underscore the importance of studying cerebellar nuclei cell function in cerebellar disease, as lack of changes in Purkinje cell firing patterns can mask disease-causing firing patterns in these cerebellar output cells.

## Introduction

The cerebellum is involved in many different movement disorders, including ataxia, dystonia, and tremor.^1^ Previous studies have found that the presentation of cerebellar disease is defined by different patterns of spike activity in cerebellar nuclei cells that bridge the cerebellar circuitry to other parts of the motor network.^2^ This suggests that spike patterns can be used as a therapeutic target or diagnostic biomarker for diverse cerebellar disorders.^3^ However, in most cerebellar disorders, cerebellar nuclei neurons are not the instigators of disease and the perturbations that cause motor impairments either impacts neurons more broadly, or predominantly affects cells upstream from the nuclei cells, cerebellar Purkinje cells.^4^

Purkinje cells are situated at the cerebellar cortex and are more readily accessible for a functional read-out than neurons in the deeper situated cerebellar nuclei.^5,6^ This makes Purkinje cells a more optimal target to measure spike signatures as biomarkers for cerebellar movement disorders but the ability to use Purkinje cell spike patterns as a read-out of cerebellar disease presentation relies on spike activity in Purkinje cells to reliably predict nuclei cell spike activity.

Purkinje cells make strongly converging, GABAergic synapses onto cerebellar nuclei cells.^6^ Based on this anatomical connectivity, it is expected that Purkinje cell activity inhibits nuclei cells (Figure 1A). Indeed, previous studies have shown that acute activation of Purkinje cell spiking activity reliably inhibits cerebellar nuclei neurons (Figure 1B).^2,7–9^ However, the effect of more prolonged or dynamically regulated changes in Purkinje cell activity is more ambiguous.^10–15^ Especially in mouse models for cerebellar disorders, like ataxia, dystonia, and tremor, there can be different relationships between Purkinje and nuclei neuron spike patterns: in a mouse model that exhibits ataxic, dystonic, and tremor symptoms (*Car8^wdl/wdl^*) both the instantaneous firing rate in Purkinje and nuclei cells is elevated; ^16^ in a different dystonia model the Purkinje cell firing rate is unchanged while a decrease in nuclei cell firing rate was observed;^17^; and in a tremor model the Purkinje cell the firing rate was decreased without a change in nuclei cell firing rate.^18^ These findings underscore the importance of gaining a better understanding of the relationship between spiking activity in Purkinje and nuclei cells in mouse models of cerebellar disease.

**Figure 1.**
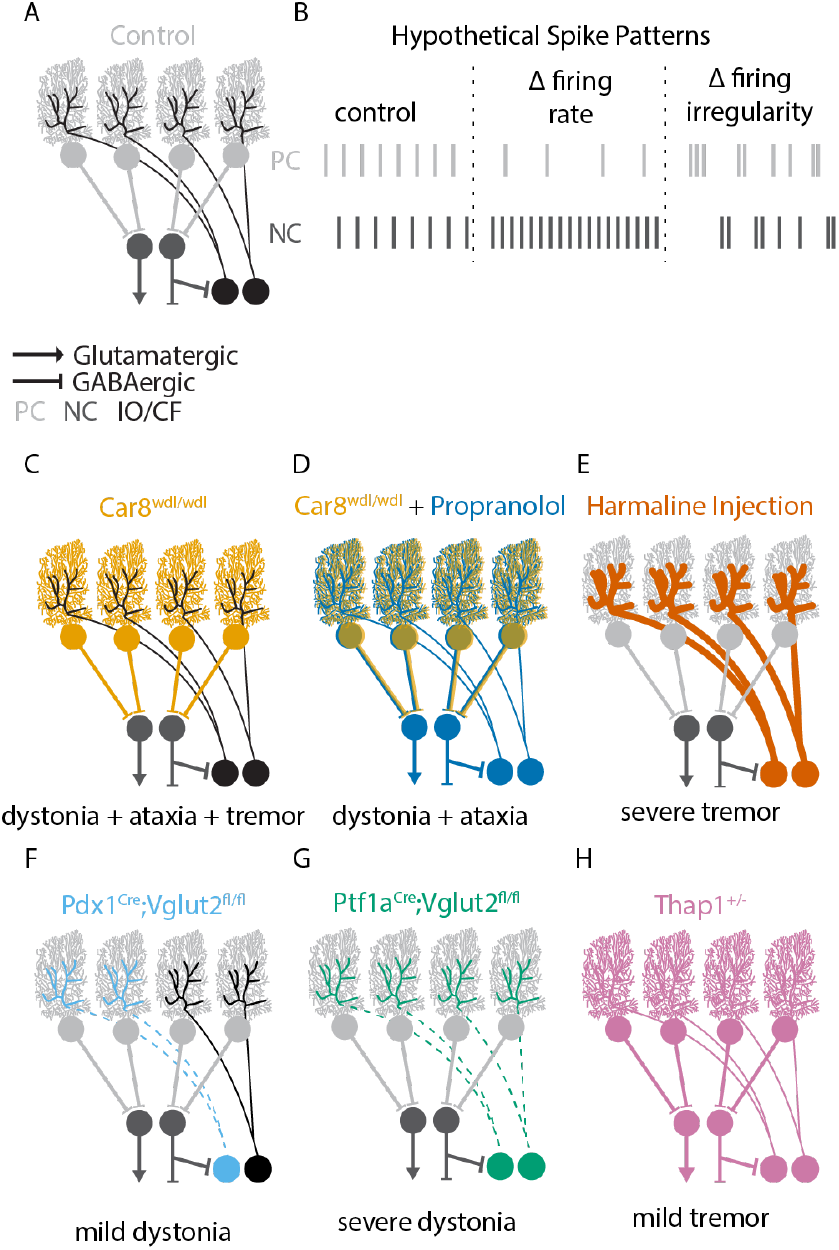
Hypothetical effects on spike patterns and affected cell types in mouse models for disease. **A** Schematic representation of a healthy cerebellar circuitry. Purkinje cells (light grey) form inhibitory GABAergic synapses on cerebellar nuclei cells (dark grey). Climbing fibers (black) from the inferior olive neurons provide excitatory glutamatergic input to Purkinje cells. **B** Examples of expected spike patterns. When Purkinje cells (light gray) have a lower spike rate than in control animals, nuclei cells (dark gray) are expected to have a high spike rate. When Purkinje cells have a high irregularity, nuclei cells should also have a high irregularity. **C-H** Schematic representation of disrupted cerebellar circuitry in mouse models for cerebellar movement disorders. The phenotype each model presents is listed below the respective schematic. Colored lines represent the cell type primarily affected; dashed lines represent no neurotransmission. Thick lines represent a pharmacologically induced mouse model, while thin lines indicate genetic mouse models.

In this study, we investigate the relationship between Purkinje and nuclei cell spike activity in a database of single-cell, *in vivo* electrophysiology recordings in healthy control mice and mouse models of cerebellar disease. Spike patterns can be described by parameters measuring spike rate (how fast action potentials are occurring) and spike irregularity (how regular the action potentials are occurring).^19^ Based on the inhibitory connection between Purkinje and nuclei cells, changes in spike rate and irregularity in Purkinje cells are likely to have different effects on downstream nuclei cells: (1) a negative correlation between Purkinje and nuclei cell spike rate; and (2) a positive correlation between Purkinje and nuclei cell spike irregularity (Figure 1B).

For our analyses, we use a dataset of in vivo, single-cell recordings from Purkinje and nuclei cells from mouse models of cerebellar disease (Figure 1C-H).^2^ The electrophysiological recordings are randomly sampled and asynchronous recordings from Purkinje and nuclei cells in the same mediolateral plane (paravermis and interposed nucleus) that are likely functioning in the same module.^20^ The nature of our data does not allow us to make predictions about direct anatomical connectivity between pairs of recordings but rather ask the question whether changes in population Purkinje cell spike rate and pattern can be used to make predictions about population nuclei cell spike rate and patten.

Our analyses include recordings from six mouse models that include single-gene genetic mouse models (*Car8^wdl/wdl^* and *^Thap1+/-^*),^16,21^ pharmacological models (propranolol and harmaline),^16,18^ and circuit-based models of selective elimination of neurotransmission in subsets of cerebellar neurons. (*Pdx1^Cre^;Vglut2^fl/fl^* and *Ptf1a^Cre^;Vglut2^fl/fl^*).^17^ Figure 1C-F summarizes which cells in the cerebellar circuit are affected by pharmacological or genetic manipulations. By using this combination of mouse models, we aim to investigate whether there are reliable relationships between spontaneously occurring (as opposed to artificially induced) changes in Purkinje and nuclei cell spike patterns.

## Results

### Range of Purkinje cell spike patterns observed in the mouse models

First, we tested whether Purkinje cell spike patterns were altered in our mouse models compared to control mice. We focused our analyses on six parameters that together richly describe spike rate (firing rate, mean instantaneous firing rate, %ISI > 25ms) and irregularity (CV, CV2, Skewness) (see methods).

We found that the firing rate was significantly reduced in two mouse models (*Car8^wdl/wdl^ +* propranolol and harmaline injected mice, Figure 2A). Mean instantaneous firing rate was signification reduced in only one mouse model (*Car8^wdl/wdl^ +* propranolol, Figure 2B). %ISI > 25 ms was increased in two mouse models (*Car8^wdl/wdl^ +* propranolol and harmaline injected mice, Figure 2C). CV was increased in three mouse models (*Car8^wdl/wdl^, Car8^wdl/wdl^ +* propranolol, and harmaline injected mice, Figure 2D). CV2 was decreased in two mouse models (*Car8^wdl/wdl^* and *^Car8wdl/wdl^ +* propranolol mice, Figure 2E) and increased in one mouse model (harmaline injected mice, Figure 2E). Finally, skewness was increased in three mouse models (*Car8^wdl/wdl^, Car8^wdl/wdl^ +* propranolol and harmaline injected mice, Figure 2F). Together, these mouse models present a range of changes in Purkinje cell spike rate and spike regularity which will allow us to investigate how these changes propagate in the cerebellar nuclei cells.

**Figure 2.**
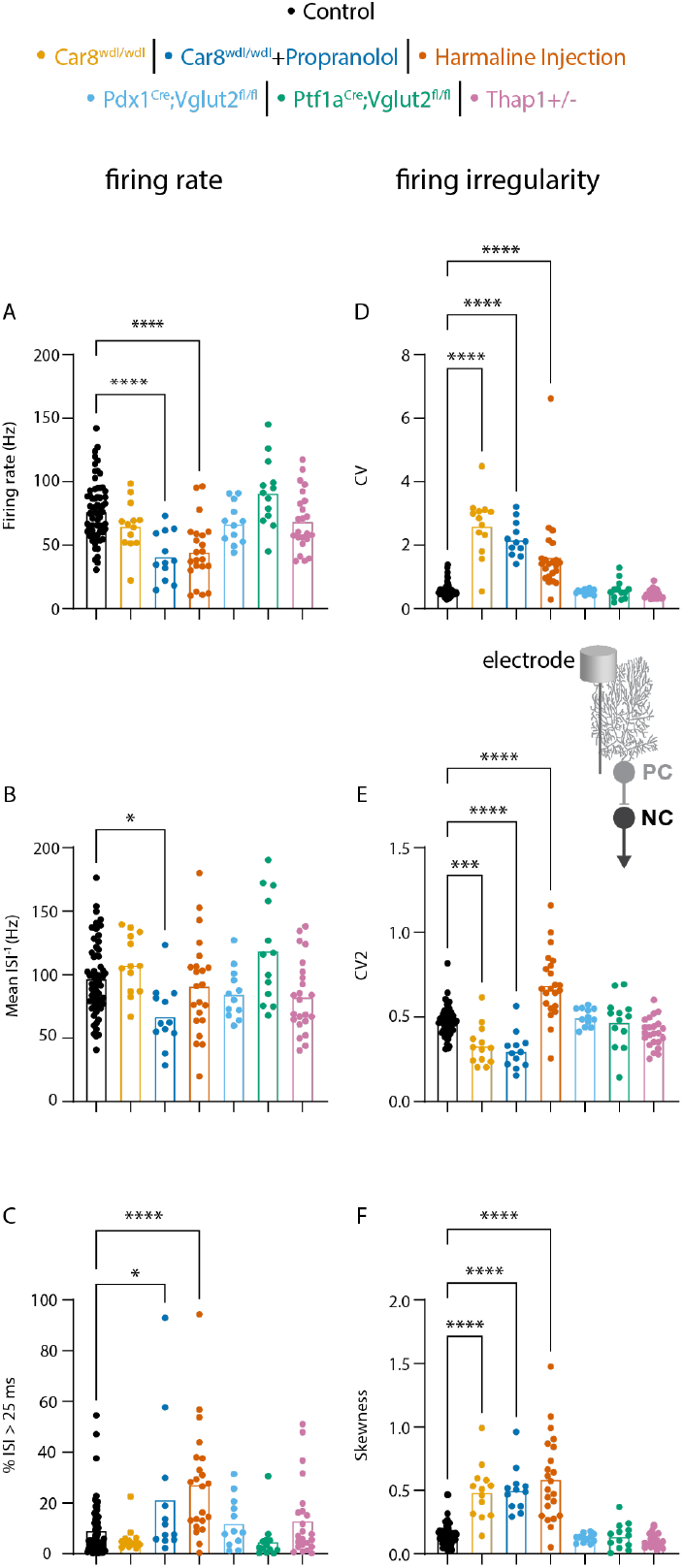
Changes in Purkinje cell spike rate and irregularity in mouse models for disease. ANOVA results for parameters relating to firing rate (A-C, left column) and firing irregularity (D-F, right column) are shown A firing rate B mean ISI-1 C percent of interspike intervals (ISI) greater than 25 milliseconds (%ISI > 25 ms) D coefficient of variance (CV) E CV2 F skewness. A representative schematic of where the electrophysiology recordings were collected is included above panel E. Here, the electrode is located at the Purkinje cell to record its spike activity. * = p-value < 0.05 | ***= p-value < 0.001 | ****= p-value < 0.0001

### Range of nuclei cell spike patterns observed in the mouse models

Next, we tested whether nuclei cell spike patterns were altered in our mouse models. We focused on the same six parameters as for the Purkinje cell spike patterns. We found that firing rate was reduced in three mouse models (*Car8^wdl/wdl^ +* propranolol, *Pdx1^Cre^;Vglut2^fl/fl^*, and *Ptf1a^Cre^;Vglut2^fl/fl^* mice, Figure 3A). Mean instantaneous firing rate was increased in two mouse models (*Car8^wdl/wdl^* and harmaline injected mice, Figure 3B). %ISI > 25ms was increased in three mouse models (*Car8^wdl/wdl^ +* propranolol, *Pdx1^Cre^;Vglut2^fl/fl^*, and *Ptf1a^Cre^;Vglut2^fl/fl^* mice, Figure 3C). CV was increased in four mouse models (*Car8^wdl/wdl^, Car8^wdl/wdl^ +* propranolol, harmaline injected, and *Ptf1a^Cre^;Vglut2^fl/fl^* mice, Figure 3D). CV2 was increased in two mouse models (harmaline injected, and *Ptf1a^Cre^;Vglut2^fl/fl^* mice, Figure 3E). Finally, skewness was increased in four mouse models (*Car8^wdl/wdl^, Car8^wdl/wdl^ +* propranolol, harmaline injected, and *Ptf1a^Cre^;Vglut2^fl/fl^* mice, Figure 3F). These data show that spike rate and spike regularity span a range in the mouse models.

**Figure 3.**
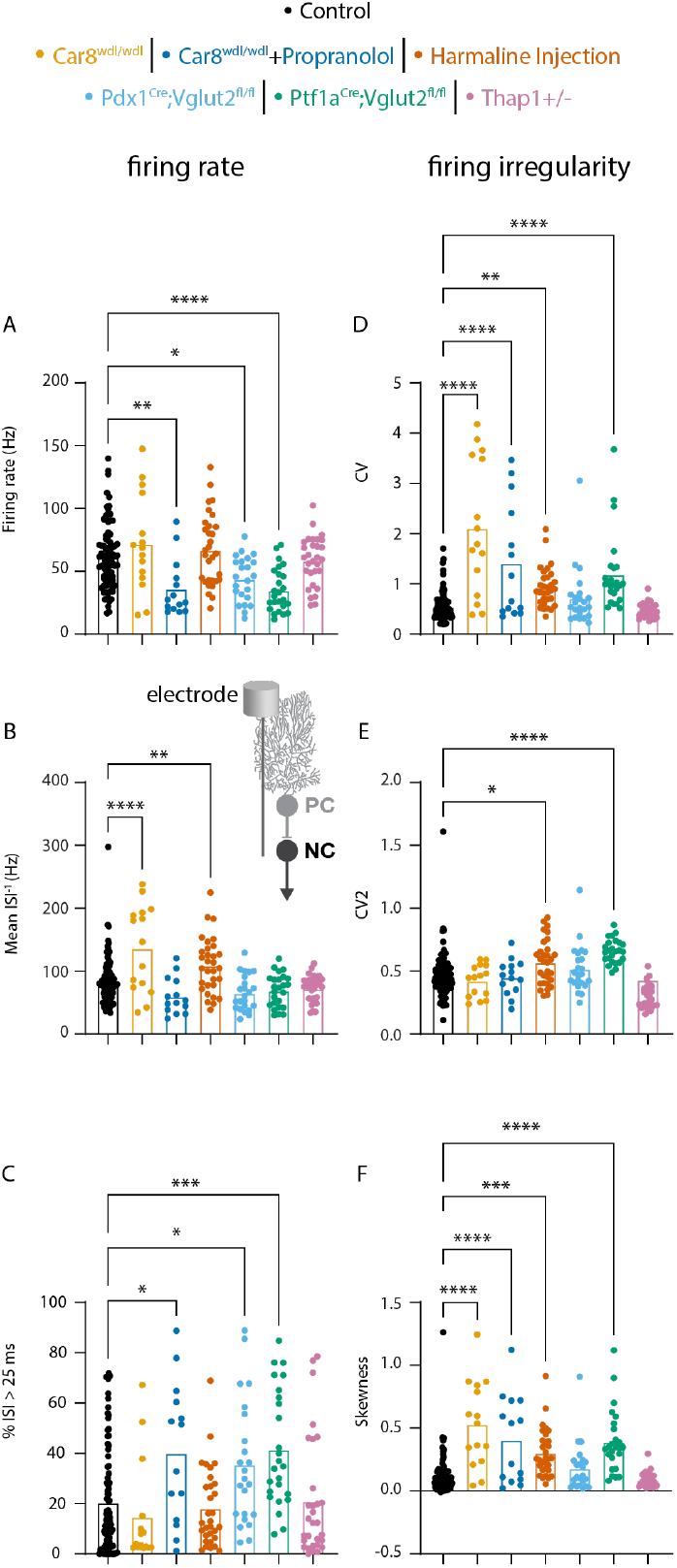
Changes in nuclei cell spike rate and irregularity in mouse models for disease. ANOVA results for parameters relating to firing rate (A-C, left column) and firing irregularity (D-F, right column) **A** firing rate **B** mean ISI^-1^ **C** percent of ISI greater than 25 ms (%ISI > 25 ms) **D** coefficient of variation (CV) **E** CV2 **F** skewness. A representative schematic of how the electrophysiology recording was collected is included between panels B and E. Here, the electrode is located at the nuclei cell to record its spike activity. * = p-value <0.05 | **= p-value < 0.01 | ***= p-value < 0.001 | ****= p-value < 0.0001

We also summarized the p-values for the one-way ANOVA post-hoc analyses comparing the Purkinje and nuclei cell spike patterns in the six mouse models to the control recordings to assess whether changes in Purkinje cell spike parameters caused reliable changes in nuclei cell parameters. Surprisingly, a statistically significant change in Purkinje cell spike rate or irregularity was not always paired with a statistically significant change in nuclei cell spike rate or irregularity. For example, a significant increase in firing rate was observed in the Purkinje cells of harmaline treated animals without a change in nuclei cell firing rate and, conversely, changes in nuclei cell firing rates were observed in *Pdx1^Cre^;Vglut2^fl/fl^*, and *Ptf1a^Cre^;Vglut2^fl/fl^* mice without changes in Purkinje cell firing rates (Table 1). These data show that statistically significant changes in Purkinje cell firing rate are neither necessary nor sufficient to induce statistically significant changes in nuclei cell firing rates.

**Table 1.**
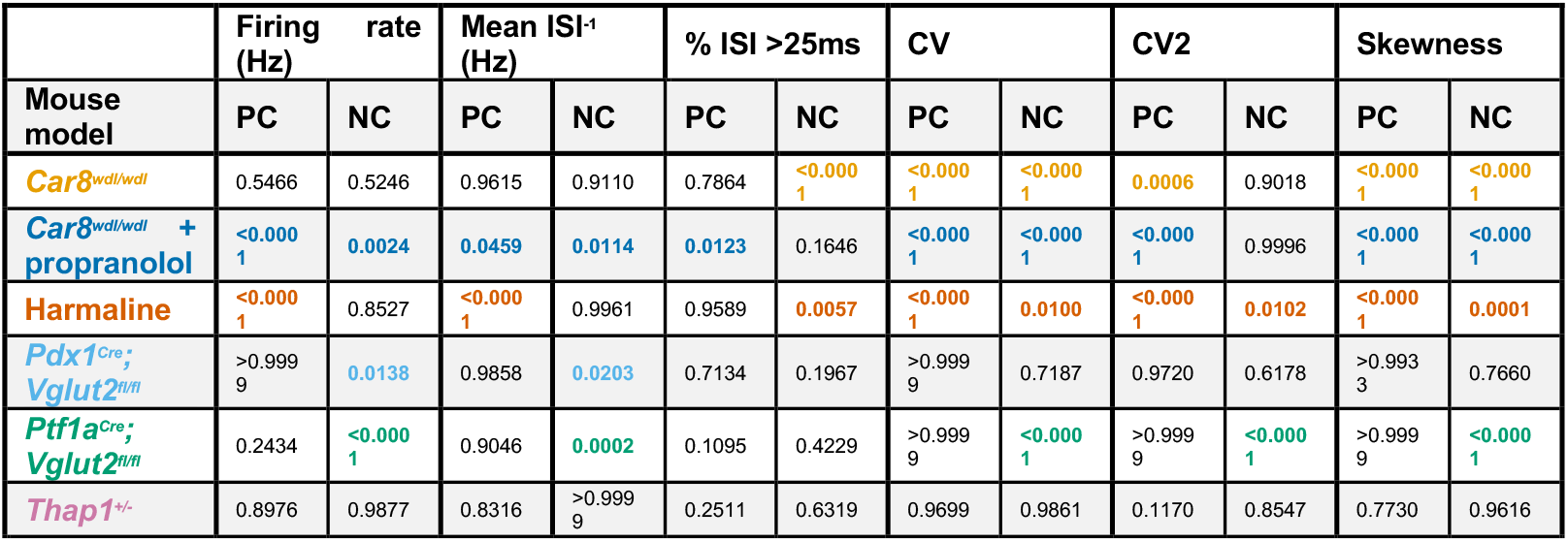
One-way ANOVA results for spike pattern differences between control and experimental mouse models. Listed are the p-values for each parameter and cell type in the different mouse models. The average of each parameter for Purkinje cells (PC) and cerebellar nuclei cells (NC) were compared to the control group for each cell type results with a Tukey post-hoc analyses for multiple comparisons. **P-value < 0.05** is considered a significant difference between means.

In addition, we also observed that changes in Purkinje cell irregularities did not cause a reliable change in nuclei cell irregularities. For example, statistically significant changes in nuclei cell CV and skewness were observed without changes in Purkinje cell irregularities in *Ptf1a^Cre^;Vglut2^fl/fl^* mice. This suggests that changes in Purkinje cell spike irregularity are not necessary to drive changes in nuclei cell spike irregularities.

### Purkinje and nuclei cell spike patterns are not correlated

The lack in co-occurring changes in Purkinje and nuclei cell spike pattern parameters may be due to insufficient power to find changes between the groups, while trends towards a certain direction may be present. Furthermore, the comparison of statistically significant effects does not ensure that the observed effects were in the same direction. We therefore also performed a linear regression analysis on the mean Purkinje and nuclei cell spike parameters (Figure 4).

**Figure 4.**
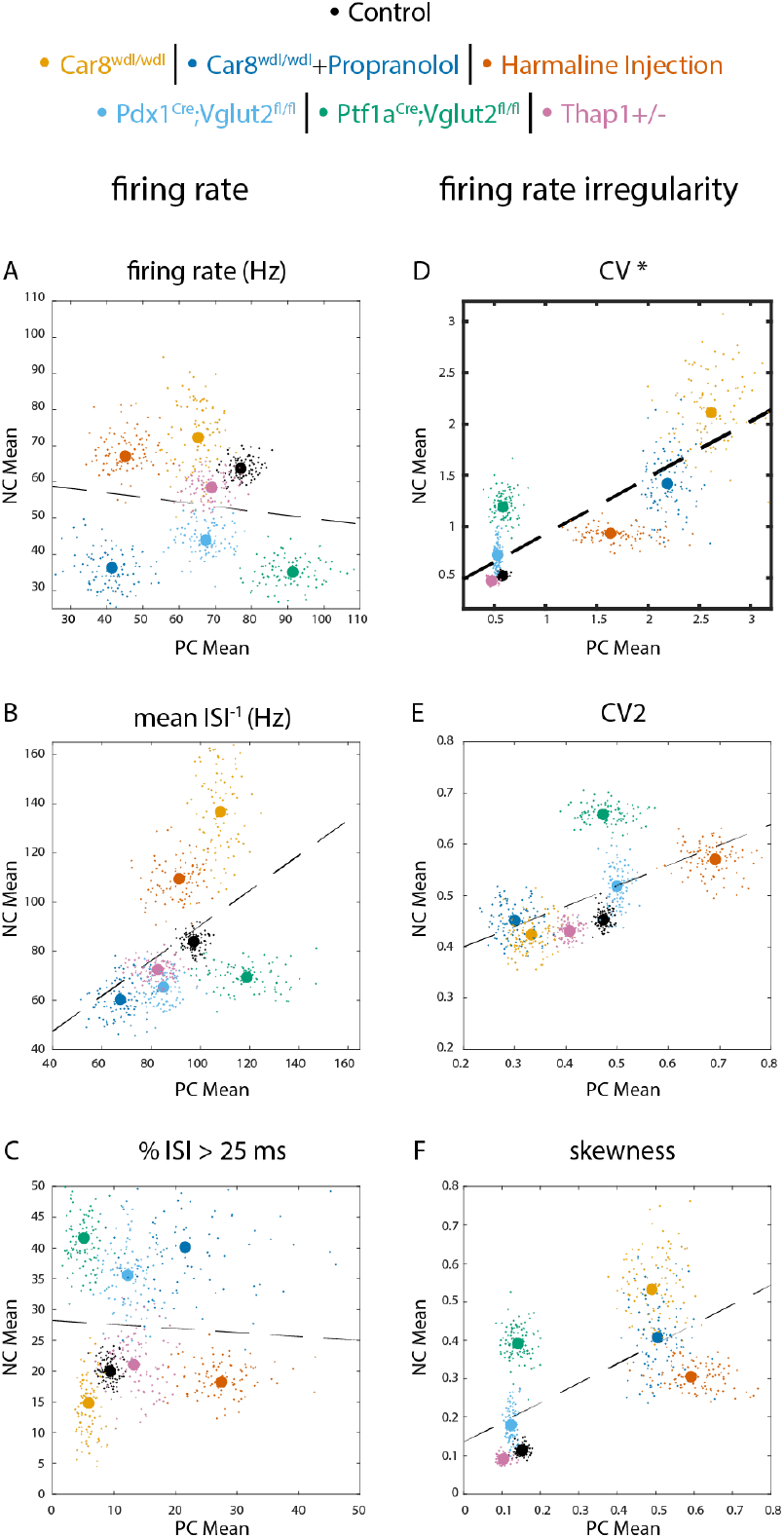
Correlation between Purkinje and nuclei cell spike rate and irregularity in mouse models for disease. Linear regression analysis and correlation plots reveal no significant relationship between Purkinje cell and nuclei cell firing rate (A). There is a positive correlation between Purkinje cell and nuclei cell irregularity (D). Correlation plots for parameters measuring firing rate are in the left column (A-C), and parameters measuring irregularity are in the right column (D-F) **A** firing rate **B** mean ISI^-1^ **C** percent of ISI greater than 25 ms (%ISI > 25 ms) **D** CV **E** CV2 **F** skewness. Dashed line in each correlation plot represents the line of best fit. Each large dot represents the true observed sample mean whereas the small dots represent a bootstrapped sample mean. **\* = p-value <0.05**

We expected to find a negative correlation between Purkinje and nuclei cell spike rate parameters and positive correlation between Purkinje and nuclei cell spike irregularity parameters. However, we found no correlation between Purkinje and nuclei cell spike rate parameters (firing rate, Figure 3A; mean instantaneous firing rate, Figure 3B; %ISI > 25ms, Figure 3C). We did find a positive correlation between Purkinje and nuclei cell spike irregularity parameter as measured by the covariance (CV, Figure 3D). However, other measures for irregularity did not show a statistically significant correlation (CV2, Figure 3E; Skewness, Figure 3F).

We also performed bootstrapped analyses to assess whether the (lack of) correlations between Purkinje and nuclei cell parameters were influenced by our specific samples or stable to changes in sampling from our dataset (Table 2). Fewer than 5% (below statistical chance) bootstrapped analyses returned statistically significant correlations between Purkinje and nuclei cell for all spike rate parameters and CV2 and skewness. However, nearly four-fifths (79%) of the bootstrapped analyses showed a significant correlation between Purkinje and nuclei cell CV. Together, these results show that changes in Purkinje cell spike CV are correlated with changes in nuclei cell spike CV but that changes in Purkinje cell spike rate are not correlated with changes in nuclei cell rate.

**Table 2.**
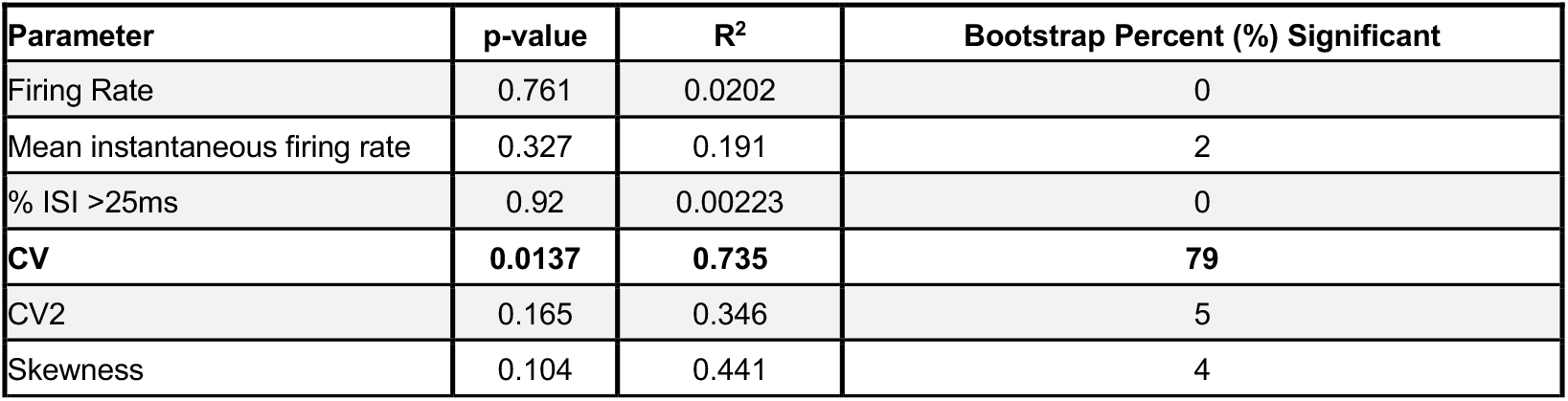
Linear regression analysis and bootstrapping results show no significant relationship between Purkinje cell and nuclei cell firing rate parameters. After completing the linear regression analysis 100 times, bootstrapping further supports no relationship between Purkinje cell and nuclei cell firing rate because little to no data points are statistically significant. The only firing irregularity parameter with a significant relationship between cell types is CV. Bootstrapping shows 79% of the data points are statistically significant for CV. The other firing irregularity parameters are not statistically significant. **P-value < 0.05**

### Nuclei cell spike patterns in mouse models of silenced or degenerating Purkinje cells

Because our findings that there was no relationship between Purkinje and nuclei cell spike regularity were against the hypothesis that a change in Purkinje cell spike rate should have an opposite effect on nuclei cell spike rate, we also tested the effects of a reduction in Purkinje cell (spikes) on nuclei cell spike patterns. In this analysis, we included three additional mouse models: *L7^Cre^;Vgat^fl/fl^* mice lack neurotransmission in from Purkinje cells, preventing Purkinje cells to influence nuclei cell spike patterns altogether (Figure 5B); *L7^Cre^;Ank1^fl/fl^* mice have severe Purkinje cell neurodegeneration due to Purkinje-cell-specific deletion of the *Ank1* gene (Figure 5C); and *Sca1^154Q/+^* mice also have Purkinje cell degeneration due a broad expression of a poly-Q knock-in the *Sca1* gene (Figure 5D). *Sca1^154Q/+^* mice were recorded earlier in disease progression than *L7^Cre^;Ank1^fl/fl^* mice.

**Figure 5.**
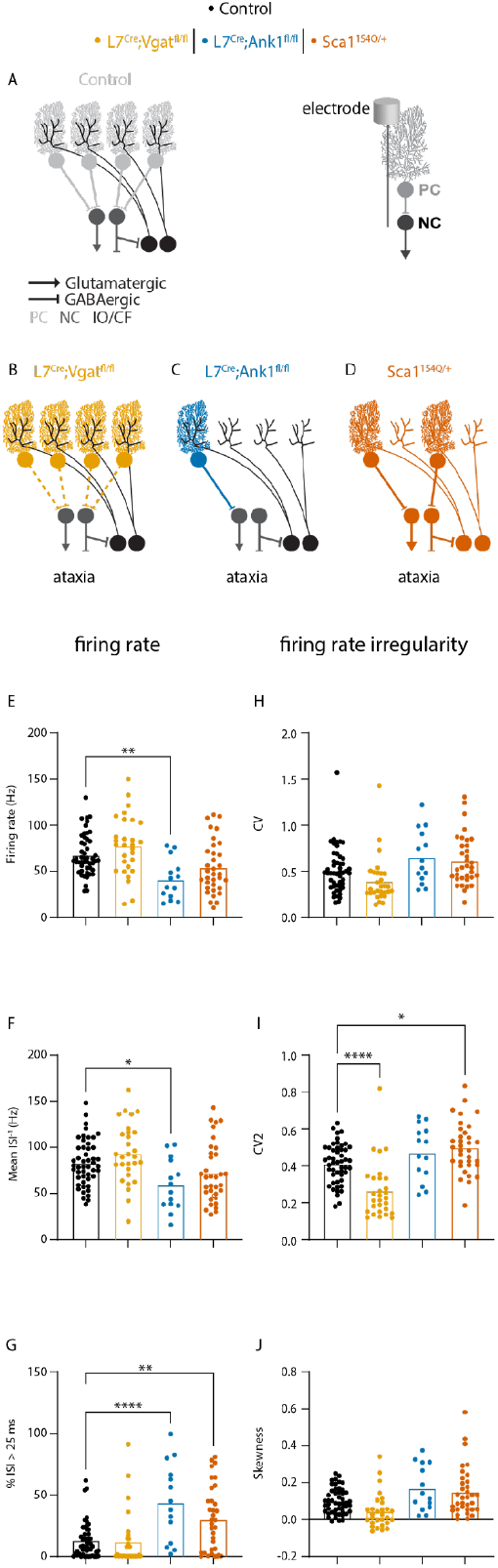
Nuclei cell spike rate is not increased in mouse models of reduced Purkinje cell output. Lack of Purkinje cell signaling does not increase cerebellar nuclei cell firing rate and has differential effects of irregularity **A-D** Schematic representation of disrupted cerebellar circuitry in mouse models for cerebellar movement disorders. These models only had impaired Purkinje cells. Colored lines represent the cell type primarily affected; dashed lines represent no neurotransmission. A representative schematic of how the electrophysiology recording was collected is included between panels B and E. Here, the electrode is located at the nuclei cell to record its spike activity. **A** schematic of control, healthy cerebellar circuitry **B** model of no Purkinje cell neurotransmission to nuclei cells **C** model of almost all Purkinje cells are missing **D** model of half of the Purkinje cells are missing **E-J** Graphs of ANOVA results for parameters relating to firing rate (left column) and firing irregularity (right column). **E** firing rate **F** mean ISI^-1^ **G** percent of ISI greater than 25 ms (%ISI > 25 ms) **H** CV **I** CV2 **J** skewness. A representative schematic of how the electrophysiology recording was collected is included next to panel A. Here, the electrode is located at the nuclei cell to record its spike activity. * = p-value < 0.05 | ** = p-value < 0.01 | ****= p-value < 0.0001

In all three ataxia mouse models, we expected to see an increase in nuclei cell firing rate, whether due to reduced inhibition from all Purkinje cells (*L7^Cre^;Vgat^fl/fl^* mice) or Purkinje cell degeneration. Nevertheless, we did not see this effect and in *L7^Cre^;Ank1^fl/fl^* mice we even observed the opposite effect of a decreased in firing rate (Figure 5F) and mean firing rate (Figure 5G). Furthermore, in *Sca1^154Q/+^* and *L7^Cre^;Ank1^fl/fl^* mice we observed a larger proportion of relatively long interspike intervals (% ISI >25ms), or low instantaneous firing rate. Together, these findings challenge the hypothesis that reduced Purkinje cell firing rates cause an increase in nuclei cell firing rates.

We also investigated how reduced Purkinje cell spikes would influence nuclei firing irregularity. We did not observe changes in global changes in nuclei cell firing irregularity (CV, Figure 5H, or skewness, Figure 5J). However, the instantaneous firing rate irregularity was significantly decreased in *L7^Cre^;Vgat^fl/fl^* mice but increased in *Sca1^154Q/+^* mice. These findings suggest that reduced Purkinje cell output has variable effects on nuclei cell firing irregularities.

## Discussion

In this paper we tested whether changes in Purkinje cell spike patterns were predictive of changes in downstream nuclei cell spike patterns. We did not observe that changes in Purkinje cell spike rate parameters were related to nuclei cell spike parameters in mouse models wherein Purkinje cell spike rates were changed (Figure 4, Table 2) or Purkinje cell spike rates were reduced (Figure 5). We did see a correlation between Purkinje and nuclei cell firing irregularity (as measured by CV) in mouse models for cerebellar disease (Figure 4, Table 2), but also found that Purkinje cell irregularity was not necessary to observe changes in nuclei cell spike changes (Figure 4, Table 1). Together, these data challenge the hypothesis that population Purkinje cell firing patterns can be used to predict downstream firing patterns.

The Purkinje and nuclei cells were recorded during the same recording session, in the same mediolateral plane, and the same mice, making it likely that we recorded neurons in the same cerebellar module.^20,22^ However, because all cells were recorded individually, we cannot confirm that our datasets reflect anatomically connected pairs of Purkinje and nuclei cells. Finding connected pairs of Purkinje and nuclei cells is technically challenging even when Purkinje and nuclei cell layers are recorded at the same time due to the convergence of Purkinje cells onto nuclei cells (∼40 Purkinje cells connect one nuclei cell) from multiple rostro-caudal planes and the synchrony of neighboring Purkinje cells.^10,12,13,23– 25^ Even though our data does not confirm Purkinje to nuclei cell connected pairs it does provide insight of the ability to predict nuclei cell output patterns based on population Purkinje cell activity. The lack of a strong relationship between Purkinje to nuclei cell firing patterns can be caused by several factors. First, nuclei cells have intrinsic abilities to generate action potentials ^26,27^ and their membrane potential adapts quickly to sustained changes in inhibitory inputs.^28^ These properties explain the tonic firing patterns in nuclei cells in the control condition during which they receive tonic inhibitory, GABAergic, inputs from ∼40 Purkinje cells firing at ∼70 Hz. This quick adaptation to changes in inhibitory tone may also protect against excitotoxicity in situations where Purkinje cell inputs are reduced, such as neurodegenerative conditions,^14,29^ or even during anesthesia when Purkinje cell firing rates drop to ∼35Hz in mice.^30^ In the disease conditions we test here, nuclei cells’ quickly adapting properties may explain the lack of relationship between Purkinje and nuclei cell firing rates. These adaptive properties, however, would not be quick enough to average out rapid fluctuations in Purkinje cell spike rates, or firing irregularity. This explains why changes in Purkinje cell firing irregularity cause reliable changes in nuclei cell firing irregularity, as measured by CV.

A second reason for a lack between Purkinje and nuclei cell firing patterns can be developmental compensation. In two of our disease models specifically, the *Pdx1^Cre^;Vglut2^fl/fl^*, and *Ptf1a^Cre^;Vglut2^fl/fl^* mice, the circuit changes resulting in mild and severe dystonia phenotypes are instigated during development. In the *Ptf1a^Cre^;Vglut2^fl/fl^* mice^17^, Purkinje cells show changes in their firing patterns that are normalized by adulthood (the dataset used for the analyses in this paper), whereas nuclei cell firing patterns continue to be abnormal throughout life. This suggests that Purkinje cell firing patterns may have instructive properties during the developmental period that define the nuclei cell’s firing patterns for life. These may be anatomical changes in the Purkinje to nuclei cell synaptic connectivity or membrane properties in the downstream nuclei cells. Regardless, changes in these developmental processes may explain why changes in Purkinje cell firing patterns in adult mice are not necessary for changes in nuclei cell firing patterns. This is an important consideration when using Purkinje cell firing patterns as a clinical indication because using Purkinje cells as a read-out of disease would mask the downstream abnormal firing patterns. Likewise, normalizing Purkinje cell firing patterns in adulthood may be insufficient to rescue the changes in circuit function that cause disease.

A final consideration for the lack of a relationship between Purkinje and nuclei firing patterns may be that nuclei neurons integrate information from ∼40 Purkinje cells^13^. Nuclei firing patterns may be influenced based on the synchrony and cross-correlations between these synaptic inputs, which information from single cell recordings do not provide ^25,31^. Multi-unit recordings may be necessary to collect sufficient information from Purkinje cells to reliably predict how they influence nuclei cell firing patterns in disease states.

In conclusion, firing patterns of a random sampled population of Purkinje cells cannot reliably predict the firing patterns of nuclei cells, which form the output from the cerebellar circuitry. This suggests that Purkinje cell firing patterns do not hold the same information that could be used as a biomarker for disease state as nuclei cell firing patterns have. Future studies are necessary to further disentangle the complicated dynamics through which populations of Purkinje cells influence nuclei cell firing patterns and how these dynamics are shaped in cerebellar disease and movement disorders.

## Methods

### Animals

Data in this paper was reanalyzed from previous publications. Detailed description of mouse lines can be found in the following citations: *Car8^wdl/wdl^* (+ propranolol)^16^; harmaline injection^18^; *Pdx1*^*Cre*^;*Vglut2*^*fl/fl2*^; *Ptf1a*^*Cre*^;Vglut2^*fl/fl*17^; *Thap1*^+/-21^; *L7^Cre^;Vgat^fl/fl^* ^18^; *L7*^*Cre*^;Ank1^*fl/fl*29^; *Sca1^154Q/+^* ^2^.

### Electrophysiological spike recordings

This study includes reanalyzed data from previously published work^2,16–18,29^. In vivo electrophysiological recordings following the experimental protocols outlined in our earlier studies. Briefly, mice were positioned on a rotating foam wheel with their heads stabilized using implanted headplates secured to the recording apparatus. The antibiotic ointment was carefully removed from the recording chamber and replaced with sterile physiological saline, unless stated otherwise. We used Bregma as a reference point to determine the coordinates for electrode insertion into the cerebellum. Recordings were performed using tungsten electrodes (2–8 MΩ, Thomas Recording) connected to a preamplifier headstage (NPI Electronic Instruments). Electrode placement was controlled via a motorized micromanipulator (MP-255; Sutter Instrument Co). Neural signals were amplified and bandpass filtered (0.3–13 kHz) using an ELC-03XS amplifier (NPI Electronics), then digitized through a CED acquisition board and recorded using Spike2 software (CED). For our analysis of cerebellar nuclei neuron activity, we only included recordings from neurons located 2.5–3.5 mm below the surface that lacked complex spikes, which are characteristic of Purkinje cell firing.

### Spike pattern parameter calculation

We used a custom-written MATLAB code to calculate the spike pattern parameters based on inter-spike-intervals (ISI) as followed:

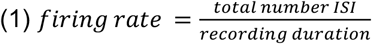

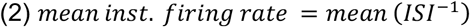

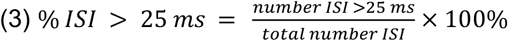

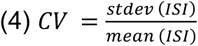

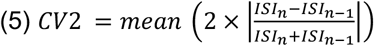

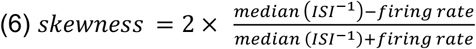

(1)*firing rate* provides the information about the average number of spikes over the duration of the recording. This parameter provides little information about firing irregularities but is sensitive to long pauses in firing patterns that are often observed in mouse models for cerebellar disease. (2)*mean instantaneous firing rate* provides a measure for firing rate that is less sensitive to long pauses and closer to the firing rate observed during bursts of activity. (3) *%ISI > 25ms* measures the proportion of inter spike intervals that are relatively long. Our previous work showed that this parameter reliably differentiated between nuclei neurons in dystonia and tremor mouse models. (4) CV measures the variability inter spike intervals and normalizes the standard deviation of inter spike intervals to the mean inter spike interval duration. This parameter is sensitive to both firing rate and firing irregularity and is often altered in mouse models of cerebellar disease. (5) CV2 measures the variability in instantaneous firing rates and is often altered in cerebellar disease models wherein the intrinsic firing properties of Purkinje cells are altered. (6) Skewness measures the difference between the average firing rate and median instantaneous firing rate. This parameter is sensitive to repeated long inter spike intervals and we previously found that it best differentiates between spike patterns in control mice versus dystonia and tremor mouse models.

### Statistical analyses

We used GraphPad Prism to analyze statistical differences in spike pattern paramaters between control and disease mouse models using a one-way ANOVA with a Tukey correction for multiple comparisons. The histograms in this manuscript were also generated using GraphPad Prism software. We used MATLAB to analyze correlation between spike pattern parameters in Purkinje and nuclei cells. We also used MATLAB to perform our bootstrap analyses and the scatterplots in this manuscript were generated using MATLAB.

## Acknowledgements

This work was supported by a Basic and Clinical Science Grant from the Dystonia Medical Research Foundation (DMRF-BCAD-2024-4) and an NIH/NINDS pathway to independence award to (R00NS130463) to MEvdH. MEvdH was also supported by start-up funds from Virginia Tech and the Red Gates Foundation. We thank members of the Van der Heijden lab and Dr. Heather Snell for feedback on the manuscript. Electrophysiology data was shared from the laboratory of Dr. Roy Sillitoe with Dr. Van der Heijden under a data transfer and usage agreement.

